# Robust, reproducible, and economical phosphopeptide enrichment using calcium titanate

**DOI:** 10.1101/457275

**Authors:** Adnan Ahmed, Vijay J. Raja, Paola Cavaliere, Noah Dephoure

## Abstract

Mass spectrometry based phosphoproteomics has revolutionized phosphoprotein analysis and enhanced our understanding of diverse and fundamental cellular processes important for human health. Because of their relative scarcity, phosphopeptides must be enriched before analysis. We demonstrate an effective and robust single-step enrichment method using an off-the-shelf preparation of calcium titanate. Our method achieves a purity and depth of analysis comparable to a widely used TiO_2_ based method at a reduced cost and effort.

---

Protein phosphorylation is the most commonly identified and best studied protein posttranslational modification. It acts as versatile chemical switch that can modulate enzyme activity, protein stability, protein interactions, and sub-cellular localization. In doing so, it regulates nearly every aspect of cellular growth, homeostasis, and proliferation^1^. Its misregulation is implicated in numerous human diseases including cancer, diabetes, and neurodegenerative disorders^2, 3^. Because of its diverse and essential regulatory roles, and because of the clinical success of protein kinase inhibitors, understanding the functions of altered phosphorylation in healthy and disease states is crucial for identifying new therapeutic targets and treatments^4,5^. The enzymes that control reversible protein phosphorylation, protein kinases and phosphatases, have been intensely studied. Since 1989, more than 1000 original journal articles with “protein kinase” or “protein phosphatase” in the title have been published each year. Despite these efforts, many of the >500 human protein kinases have no known substrates and more than a third have poorly or undefined functions^6^. We have, in all likelihood, only scratched the surface of our understanding of an immensely complex network of protein phosphorylation whose features have profound effects on human health.

With the advent of accurate quantitative multiplexing, enabled by isobaric chemical labeling,^7,8^ mass spectrometry based phosphoproteomic methods now provide a robust experimental platform for temporal and condition specific study of protein phosphorylation^9, 10^. The provided higher throughput and quantitative platform enable the broad exploration of the mechanisms and functions of protein phosphorylation in diverse conditions and across many samples. However, these experiments remain costly and time consuming. Their wider adoption will require new sample preparation and analysis methods. Here we demonstrate the efficacy of a simple and economical method for purifying phosphopeptides from complex peptide mixtures using calcium titanate (CaTiO_3_).

The success of large-scale phosphoproteomics arose from, and relies upon, methods for enriching relatively scarce phosphopeptides from total protein digests prior to mass spectrometric analysis^11, 12^. This is commonly achieved using immobilized metal affinity chromatography (IMAC) or titanium dioxide (TiO_2_)^13^. In recent years, optimized methods using commercial preparations of TiO_2_ (Titansphere) have become popular due to their reliable performance^14, 15^. However, to achieve deep coverage of the phosphoproteome it is necessary to start with large amounts of protein, as much as 10 mg or more and even larger amounts of TiO_2_ ^9^. Analyzing many samples, such as is done in TMT multiplexing, could require close to 400 mg of commercial TiO_2_ at a significant cost. In our search for more economical solutions, we were surprised to find very little data examining the relative phosphopeptide enrichment efficacy of different sources and forms of TiO_2_. Thus, we set out to perform such tests, starting with the two primary mineral sources and crystal forms of TiO_2_, rutile and anatase, and to compare them to the commonly used Titansphere particles from GL sciences. Alongside these different TiO_2_ preparations, we also included a less commonly used material, CaTiO_3_, that was reported to provide high phosphopeptide selectivity^16^. For both forms of raw TiO_2_ and CaTiO_3_, we used Jurkat cell extracts digested with lysyl endopeptidase (lysC) and trypsin to test two different ratios of enrichment substrate to peptides, 4:1 and 10:1, and two different protocols, the optimized Titansphere protocol which uses a viscous solution of 2 M lactic acid in 50% ACN followed by washes in a ACN and TFA buffer and the other with a simplified buffer system using only a 50% ACN and 1% TFA buffer. We compared all of these to the optimized protocol using Titansphere resin. All tested materials and conditions generated sizeable phosphopeptide datasets from a single 85-min LC-MS/MS analysis on an Orbitrap Fusion mass spectrometer (Fig 1, supplemental dataset 1). CaTiO_3_ outperformed both rutile and anatase TiO_2_. Rutile TiO_2_ was the only material that failed to surpass 5,000 identified phosphopeptides in at least one condition. Surprisingly, a crude preparation of −325 mesh (particle size <44 um) CaTiO_3_ at 10:1 (wt:wt) CaTiO_3_ to peptide in a TFA buffer was as effective as the optimized protocol using Titansphere, a high grade commercial preparation of spherical narrow size range particles (Fig 1).

**Fig 1.**
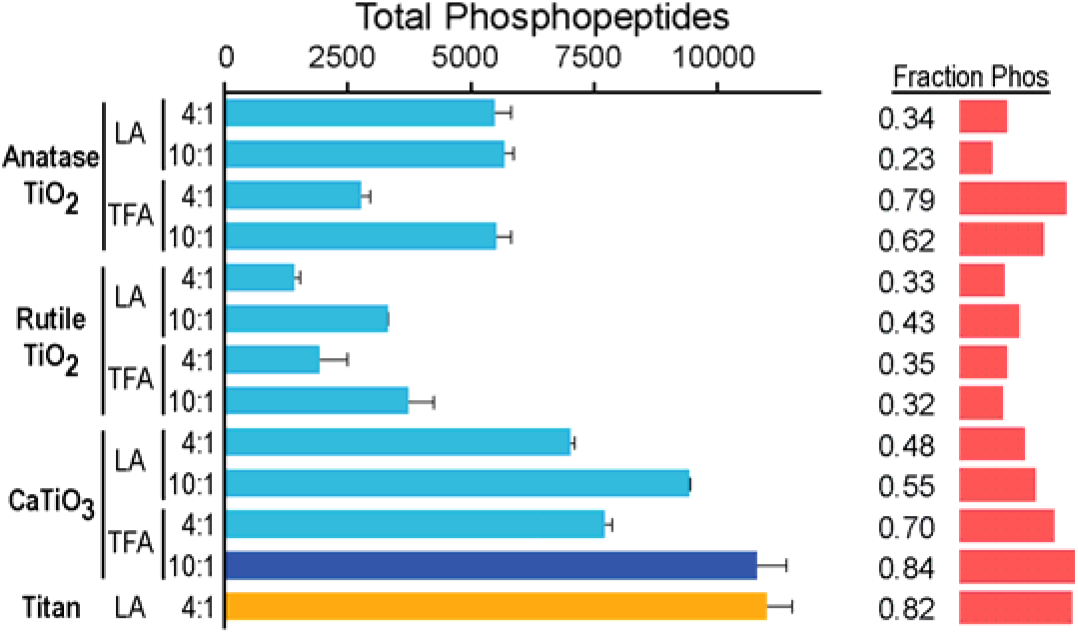
Comparing phosphopeptide enrichment materials. The average total number of phosphopeptides identified from two independent purifications using each of the indicated combinations of resin, buffer, and ratio of resin to peptides (wt:wt). LA = lactic acid, TFA = trifluoroacetic acid. Error bars indicate the standard deviation. Shown at right are the corresponding fraction of all identified spectra matched to phosphorylated peptides. All experiments used 0.5 mg aliquots of lysC and trypsin digested Jurkat lysate.

To confirm this result and further benchmark CaTiO_3_ as a phosphopeptide enrichment tool, we performed triplicate analysis from 1 mg of both Jurkat and HeLa extracts using a TFA/CaTiO_3_ method alongside the optimized Titansphere method. Both CaTiO_3_ and Titansphere consistently yielded ≥90% phosphopeptide enrichment (Fig 2a and Fig S1), and both were highly reproducible (Fig 2a and 2b, Fig S1a and S1b). CaTiO_3_ particles proved to be extremely robust and consistently performed as well or better than Titansphere. CaTiO_3_ enrichment yielded as many or more phosphorylated peptide spectral matches (PSM), unique phosphopeptides, and localized sites while providing similar phosphopeptide purity and signal intensity (described below) at a cost of ~$0.0005/mg of peptide, >2000-fold less than that for Titansphere TiO_2_.

**Fig 2.**
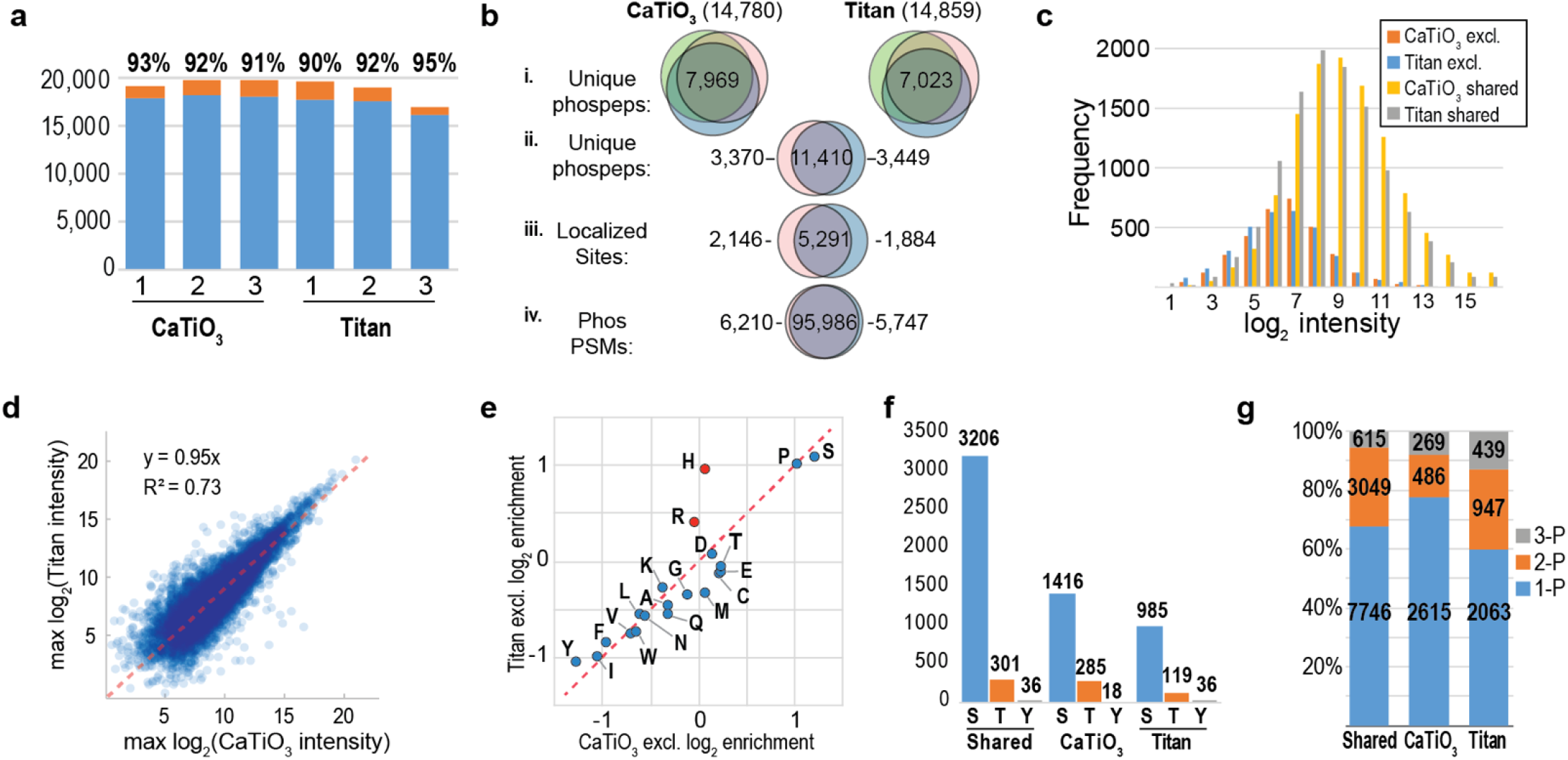
Benchmarking CaTiO_3_ phosphopeptide enrichment against Titansphere TiO_2_ enrichment in Jurkat Cell extracts. **a**, CaTiO_3_ and Titansphere provide comparable numbers of phosphopeptides with similar purity. The total number of spectra matched to phosphopeptides (blue) and non-phosphorylated (orange) peptides is shown. **b**, Overlap analysis of the unique sequences and sites identified by three trials of each method (i), overlap of unique phosphopeptides between the combined datasets for each (ii), overlap of confidently localized sites between methods (iii), and the overlap of all spectral matches between methods (iv). **c**, Commonly identified sequences are of higher intensity. The distribution of maximum log_2_ signal intensities of phosphopeptides is plotted separately for shared and exclusive (excl.) phosphopeptides identified by each method. **d**, Peptides identified by both CaTiO_3_ and Titansphere enrichment have similar intensities. For each unique sequence in common, the maximum signal-to-noise value for each method is plotted. **e**, Amino acid frequencies of CaTiO_3_ and Titansphere exclusives phosphopeptides are plotted as the log_2_ ratio of their frequencies relative to those found in a single 85 min analysis of unenriched HeLa extract peptides. **f**, The distribution of Ser, Thr, and Tyr phosphorylations among confidently localized singly phosphorylated peptides exclusive to each method. **g**, Titansphere enrichment identifies more multiply phosphorylated peptides. The distributions of singly (1-P), doubly (2-P), and triply (3-P) phosphorylated peptides found with each method independently is shown along with the number of shared sites. Similar data for HeLa derived peptides appears in Fig S1.

Analysis of the identified phosphopeptides revealed a large overlap but some clear differences. Phosphopeptide comparisons are complicated by the frequent inability to unambiguously localize the site of phosphorylation to a single amino acid. We used two different methods to make this comparison. First, we used a conservative approach in which we grouped peptides with the same amino acid sequence and the same total number of phosphorylation sites to generate the minimum number of distinct phosphopeptides present in the dataset. By this metric, we identified 18,229 unique phosphopeptide sequences from the Jurkat cell extracts, 11,410 of which (63%) were identified by both methods (Fig 2bii). The remaining unique sequences were divided almost equally between the two methods. As an alternative means of comparing the overlap of CaTiO_3_ and TiO_2_ phosphorylation datasets, we used the AScore algorithm^17^ to assess the probability of correct site assignment and examined the overlap of singly phosphorylated peptide derived sites that were localized with ≥99% probability (Ascore ≥19). We identified 9,321 confidently localized sites, 5,291 of which (57%) were found in both datasets with the remainder again being split nearly evenly between the two (Fig 2biii). Parallel experiments performed with HeLa cells generated similar results (Fig S1).

Because we identified many phosphopeptides repeatedly, we also examined the overlap considering the fraction of total spectral matches from common and unique sequences. Notably, the vast majority of matched spectra corresponded to shared sequences. Of the 102,353 PSMs identified collectively in six runs, 95,986 (94%) corresponded to the 11,410 sequences found at least once with both methods (Fig 2b iv). Shared sequences were identified by an average of 8.4 spectra. Phosphopeptides found exclusively after enrichment with either CaTiO_3_ or Titansphere were identified at a far lower rate suggesting they represent less abundant species. The number of PSMs matching sequences found exclusively in only one of the two experiments was similar - 6,210 for CaTiO_3_ and 5,747 for Titansphere. Exclusive sequences were identified by an average of 1.8 PSMs, indicating that they were usually not found in all three replicates of each method. We further inspected the observed intensity of common and exclusive sequences by looking at the distribution of precursor signal-to-noise measurements. Titansphere and CaTiO_3_ exclusive peptide intensities were on average 3-4 fold lower than commonly identified ones (Fig 2c). We then looked at the set of commonly identified peptides to see if either method generated greater signal intensities that might benefit quantitative analysis. Comparing the maximum signal-to-noise value for each method showed a strong linear correlation (R^2^=0.70) and no intensity bias (Fig 2d).

Though both methods produced a comparable quality and depth of phosphopeptide datasets, and the vast majority of identified spectra matched sequences found with both, each also generated a sizeable number of unique phosphopeptides and phosphorylation sites. To better characterize these differences, we compared a range of peptide properties including length, missed cleavages, mol wt, and hydrophobicity along with the frequency of different classes of amino acids (Table S1). Titansphere enriched peptides were slightly longer, heavier, more likely to have missed cleavages, and more basic. A closer examination of individual amino acid frequencies revealed a marked increase in the frequency of the basic residues histidine and arginine in the Titansphere exclusive phosphopeptides (Figs 2e, S1e, Table S1). The subtle but reproducible increase in peptide length and mol wt. observed with Titansphere can likely be explained by an increased affinity for basic residues whose numbers are increased on missed cleavage peptides.

By looking at confidently localized sites (Ascore ≥19), we found that the relative proportion of phosphoserine and phosphothreonine sites was nearly identical (Fig 2f). Titansphere enrichment recovered slightly more tyrosine phosphorylated sites, 72 (36 exclusive) versus 54 (18 exclusive) for CaTiO_3_. Though subtle, a similar disparity was also observed in HeLa cells (Fig S1f), where we identified 105 (45 exclusive) versus 74 (14 exclusive) localized phosphotyrosine sites. The most notable difference between the two methods was that Titansphere enrichment recovered more multiply phosphorylated sequences (Fig 2g). Each method identified a common set of 3664 multiply phosphorylated sequences, but Titansphere identified nearly twice as many doubly and triply phosphorylated sequences as CaTiO_3_ (1386 versus 755). This result was recapitulated in HeLa cells (Fig S1g). The increased length of Titansphere enriched phosphopeptides due to the higher incidence of missed cleavages may account for the observed increase in multiply phosphorylated peptides. Longer peptides were observed to have more phosphorylation sites in both cell lines and with both methods (Fig S2), and peptides with more phosphorylation sites were longer; doubly and triply modified sequences were on average three and five residues longer than singly phosphorylated peptides, respectively. These data demonstrate how subtle changes in purification methods can generate distinct phosphopeptide datasets.

We have demonstrated a remarkably robust and scalable method for generating large pools of enriched phosphopeptides for mass spectrometric analysis using widely available and affordable CaTiO_3_. In experiments in two cell lines, we showed it to be as reliable and effective as a popular method using a commercial preparation of TiO_2_. Whether used to supplant or supplement existing methods, the ease and economy of CaTiO_3_ phosphopeptide enrichment will facilitate future large-scale phosphoproteomic analyses of diverse samples.

## Methods

Online methods are provided in a separate supplementary document.

## Acknowledgements

We thank Yogindrya Vedyvas and Moonsoo Jin for providing and verifying the identify of Jurkat and HeLa cells. Noah Dephoure and Vijay Raja are supported in part by a grant from Project ALS. This work was also funded by the Dephoure laboratory startup package.

## Author Contributions

N.D. supervised all aspects of the project. A.A., V.J.R., and P.C. performed experiments. A.A., V.J.R., P.C., and N.D. performed data analysis. ND wrote the manuscript. All authors discussed the results and commented on the manuscript.

## Competing Interests

The authors declare no competing interests.

## Supplementary information

Including a single table and five supplementary data files in excel format are provided in separate documents. Two supplementary figures are included at the end of this document. Raw mass spectrometric data have been deposited in the PRIDE archive (https://www.ebi.ac.uk/pride/archive/) as dataset PXD011145.

